# Beneficial effects of prolonged 2-phenylethyl alcohol inhalation on chronic distress-induced anxio-depressive-like phenotype in female mice

**DOI:** 10.1101/2022.03.17.484716

**Authors:** Bahrie Ramadan, Lidia Cabeza, Stéphanie Cramoisy, Christophe Houdayer, Patrice Andrieu, Jean-Louis Millot, Emmanuel Haffen, Pierre-Yves Risold, Yvan Peterschmitt

**Author notes:** Corresponding authors at: Laboratoire de Recherches Intégratives en Neurosciences et Psychologie Cognitive – UR-LINC 481, 19 Rue Ambroise Paré, 25030 Besançon Cedex, 0033 3 63 08 22 18., E-mail addresses (B. Ramadan), (Y. Peterschmitt). Deceased (J-L. Millot), contributed for the study concept and design.

## Abstract

Chronic distress-induced hypothalamic-pituitary-adrenal axis deregulations have been associated with the development of neuropsychiatric disorders such as anxiety and depression. Currently available drugs treating such pathological conditions have limited efficacy and diverse side effects, revealing the need of new safer strategies. Aromatic plant-based compounds are largely used in herbal medicine due to their therapeutic properties on mood, physiology, and general well-being. The purpose of this study was to investigate the effects of 2-phenylethyl alcohol (PEA), one of the pharmacologically active constituents of rose essential oil, on chronic corticosterone (CORT)-induced behavioral and neurobiological changes in female mice. Animals followed a prolonged PEA inhalation exposure (30 min per day) for 15 consecutive days prior to behavioral evaluation with open-field, forced swim and novelty-suppressed feeding tests. CORT treatment induced an anxio-depressive-like phenotype, evidenced by a reduced locomotor activity in the open-field, and an increased latency to feed in the novelty-suppressed feeding paradigms. To elucidate the neural correlates of our behavioral results, cerebral cFos expression analysis was further performed to provide a global map of neuronal activity. The altered feeding behavior was accompanied by a significant decrease in the number of cFos-positive cells in the olfactory bulb, and altered brain connectivity as shown by cross-correlation-based network analysis. CORT-induced behavioral and neurobiological alterations were reversed by prolonged PEA inhalation, suggesting a therapeutic action that allows regulating the activity of neural circuits involved in sensory, emotional and feeding behaviors. These findings might contribute to better understand the therapeutic potential of PEA on anxio-depressive symptoms.

## 1. Introduction

Mental disorders represent a major public health concern with an important socioeconomic burden worldwide. Among them, anxiety and depression are characterized by a wide symptomatology [1] that significantly impacts patients’ quality of life. These neuropsychiatric disorders are among the most common in today’s society and present a higher prevalence among women than men [2,3]. Although nosological diagnosis considers anxiety and depression as two distinct diseases [1], clinical practice demonstrates that they often co-exist, which challenges appropriate therapeutic strategies and medical care [4,5]. Distress is considered the major external environmental factor that may trigger the emergence of anxio-depressive disorders, as it has been extensively documented (for review see [6]). Of particular interest for the present investigation, chronic distress correlates with hypothalamic-pituitary-adrenal (HPA) axis dysfunction in response to sustained elevated glucocorticoids (GC) secretion [7–9]. Even though anxiety shows a wider spectrum of HPA axis activity patterns than depression [6], these disorders can be understood as consecutive phenotypes in a physiological to pathological continuum [10], since their etiology often overlaps and relates to altered patterns of the stress-response system.

Preclinical investigation has extensively contributed to understand the relationship between chronic distress and the emergence of mood disorders, and in particular chronic GC administration in rodents has allowed to address their pathophysiology by mimicking stress-induced HPA dysfunction [11]. Indeed, sustained GC administration exerts anxiogenic-like effects on rodents’ behavior, as demonstrated in the open-field (OF) and elevated plus maze (EPM) tests, the light/dark box task and the novelty-suppressed feeding (NSF) task [12–19]. Treated animals also exhibit cognitive dysfunctions [16,18–20], including motivational deficits, suboptimal decision-making under uncertainty and suboptimal spatial working memory processing, as recently demonstrated in male C57BL/6JRj mice in our laboratory [21,22]. Most importantly, literature has reported that the pathological phenotype induced by sustained GC administration is sensitive to chronic antidepressant treatment [17, 18, 27], which supports the reliability of the model for screening the pharmacological value of new drugs [24].

Classical medications designed to treat anxio-depressive symptoms have frequently limited efficacy and diverse side effects [25–27]. In an attempt to develop safer and better-tolerated therapeutic strategies, over the past decades there has been a growing interest in the potential beneficial effects of substances derived from aromatic plants. In this context, *aromatherapy* (i.e., ‘the use of essential oils extracted from plants for the treatment of physical and psychological health’ [28]) is increasingly being considered as an alternative or therapeutic adjuvant for the improvement of patients’ mental well-being. It is therefore conceivable that diffusion of essential oils in psychiatric wards could be a convenient method to reduce problems of poor adherence to medication treatment and to improve patients’ quality of life. Even though data concerning the effectiveness of different essential oils in clinical context is still lacking, it appears that the most effective route of application is inhalation, and the physiological changes induced by aromatherapy can be sustained when inhalation is repeated [29].

Rose essential oil is one of the most commonly used plant extracts in herbal medicine [30]. For instance, beneficial effects have been reported for the use of rose essential oil during childbirth, specifically reducing maternal feelings of anxiety [31]. Similarly, anxiolytic effects have been described during labor, when rose essential oil is inhaled and used in footbaths [32]. In this concern, it has been suggested that these effects could be mediated by the action of the aromatic extract on the HPA axis, since its exposure decreases salivary cortisol levels in healthy female students subjected either to chronic or acute stressful situations [33]. Animal studies using rose odor have also evidenced its positive influence on behavior. Rose essential oil has an anti-conflict effect in ICR male mice [34], a potential anxiolytic effect in Wistar male rats [35], and in Mongolian gerbils [36]. Similar to human data, acute rose essential oil inhalation decreases GC plasma levels in rats subjected to acute restraint stress [33], and reduces adrenocorticotropic hormone plasma levels in C57BL/6J male mice exposed to predator odor [37]. Most importantly, it has been pointed out that the anxiolytic profile of rose odor strengthens after chronic inhalation in rodents [36].

Among the various chemical components of rose essential oil, 2-phenylethyl alcohol (PEA) has been suggested to be one of its main pharmacologically active constituents [38]. By itself, PEA is able to induce the anti-conflict effect observed in ICR male mice exposed to rose oil [34,38]. However, only few studies have evaluated the therapeutic effect of PEA on behavior. Specifically, our laboratory proposed an anxiolytic-like profile of acute PEA exposure in OF1 female mice, suggested by a reduction in the occurrence of anxious-like behaviors in the EPM test and subsequent decreased GC plasma levels [39]. A more recent study has reported an anxiogenic effect of acute PEA inhalation in C57BL/6 male mice tested in the OF test, but an anti-depressive effect in the tail-suspension test, thus suggesting its potential as alternative/adjuvant treatment for mood-related disorders [40].

This investigation further examines the therapeutic properties of PEA in female mice chronically treated with exogenous GC. In particular, we evaluated whether chronic inhalation of PEA could reverse the sustained GC-induced behavioral alterations and explored their underlying neurobiological mechanisms through functional network representations.

## 2. Materials and methods

### 2.1. Animals

Forty 6-to 8-week-old C57BL/6JRj female mice (*Ets Janvier Labs*, Saint-Berthevin, France) were used in this study (mean weight ± SEM (g) at the beginning of the experiments = 18.85±0.16). Mice were group housed (n=5 per cage) in Plexiglas cages and maintained in a temperature and humidity-controlled environment (22±2 °C, 55±10% respectively) under a 12 hours light/dark cycle. Animals had access to food (KlibaNafag3430PMS10, *Serlab*, CH-4303 Kaiserau, Germany) and water or treatment *ad libitum*, and were weekly handled and weighted.

All procedures were performed in accordance with the European Community Council Directive of September 22^nd^, 2010 (2010/63/UE). The experiments were approved by the Ethical Committee in Animal Experimentation from Besançon (CEBEA-58; A-25-056-2) and all efforts were made to minimize animal suffering.

### 2.2. Pharmacological treatment

Corticosterone (CORT, 4-Pregnene-11β-diol-3,20-dione-21-dione, *Sigma-Aldrich*, France) was dissolved in vehicle solution (VEH, 0.45% hydroxypropyl-β-cyclodextrin-βCD, *Roquette GmbH*, France) and administered orally through drinking water (35 μg/ml equivalent to 5 mg/kg/day; CORT group, n=20) throughout the entire experimental procedure. Control animals received vehicle solution dissolved in the drinking water (VEH group, n=20). The dose and duration of treatment were selected based on previous studies [13,21,41].

### 2.3. Chronic olfactory inhalation

A highly concentrated solution of PEA (99% C_8_H_10_O=122.17 g.mol^-1^, *Acros Organics*®, France) was used in the current study in line with our previous data [39]. Distilled water mimicking a neutral odor condition was used as control. In order to expose the animals to olfactory inhalation, the home cages with removed filter lids were introduced in a closed wooden box (length: 79, width: 40 and height: 41 cm). Cotton gauze impregnated with 239μl of PEA (or distilled H_2_O) was fixed on the top of an axial fan allowing homogenous odor diffusion within the experimental box. Chronic olfactory inhalation protocol, prior to behavioral testing, lasted 15 consecutive days (30 min per day). The duration of prolonged odor exposure was selected based on a previous study [36].

In order to evaluate the influence of chronic PEA exposure on emotional behavior and neuronal activation, mice that were previously divided into the CORT and VEH groups, were further assigned to the following experimental conditions: VEH+H_2_O, VEH+PEA, CORT+H_2_O and CORT+PEA, all n=10.

### 2.4. Behavioral assessments

Behavioral evaluations started on the 8^th^ week of VEH/CORT treatment and were conducted during the light phase of the circadian cycle in an isolated behavioral room with a dim light adjusted for each test. Behavior was video-taped for ulterior scoring. A timeline of the experimental design is presented in **Fig. 1**.

**Figure 1.**
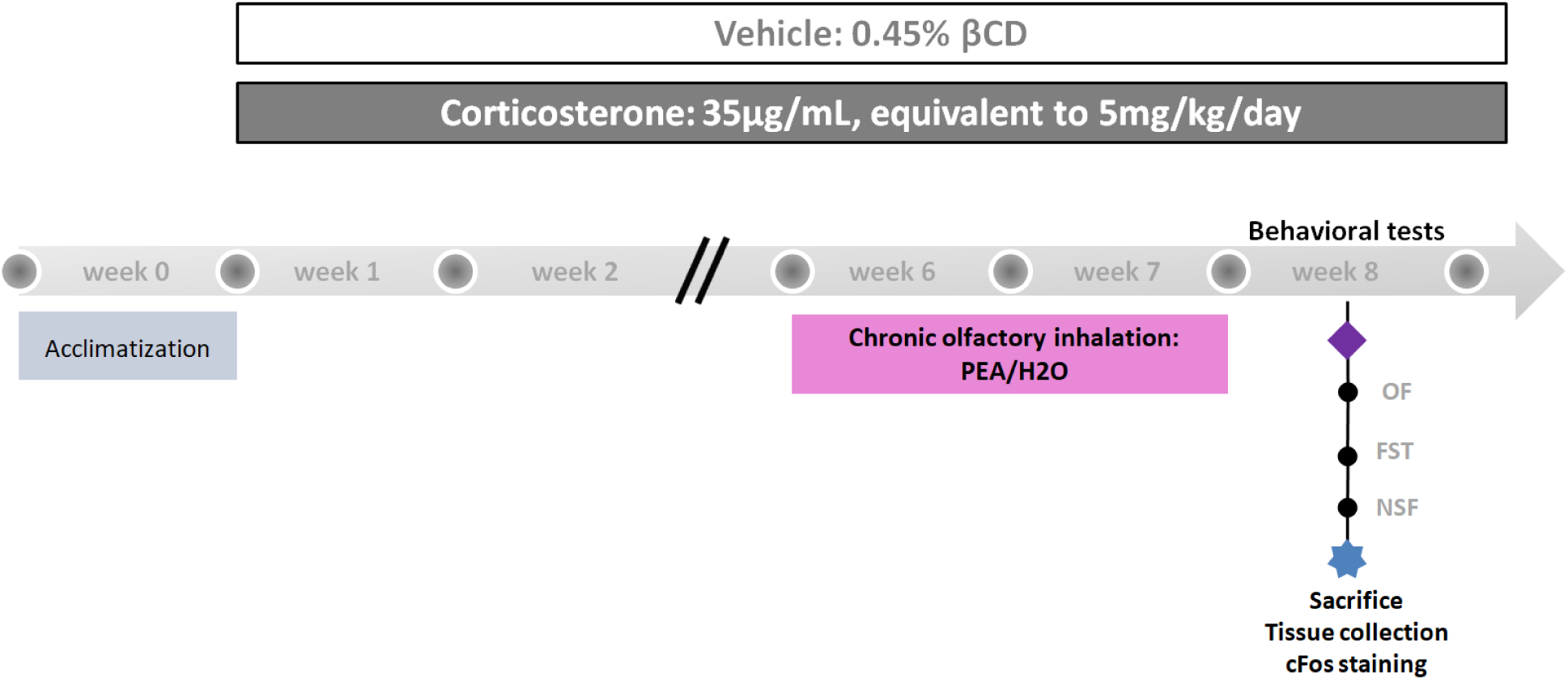
Schematic representation of the experimental design. After an acclimatization period of one week, mice were treated with corticosterone (CORT, 35 μg/mL; n=20) or vehicle (VEH, 0.45% hydroxypropil-β-cyclodextrine-βCD; n=20) for 8 weeks. Olfactory exposure to either distilled H_2_O or PEA (2-phenethyl alcohol) took place 30 min per day, for 2 weeks from the 6^th^ week of treatment onwards. The behavioral evaluation was performed during the 8^th^ week. Mire were serially tested in the open-field (OF) test, the forced swim test (FST) and the novelty-suppressed feeding (NSF) task in order to evaluate the effects of prolonged PEA inhalation on chronic CORT-induced anxio-depressive-like phenotype. Sixty-to-eighty min after the NSF evaluation, mice were sacrificed (n=5, all conditions) and their brains harvested and processed for cFos immunostaining.

#### 2.4.1. Open-Field Test

This behavioral test allows approaching anxious-like behavior as previously reported [42]. Mice (n=40) were individually placed in the center of an opaque cylindrical arena (diameter: 40, height: 48 cm), which was virtually divided into two zones: a central highly illuminated zone (diameter: 34 cm; 100 lux) and a peripheral more slightly illuminated zone (diameter: 3 cm; 60 lux). The test lasted 5 min and the time spent in the central zone was assessed as an index of anxious-like behavior. Effects on locomotor activity were evaluated based on the animals’ total distance traveled in the arena [39,43].

#### 2.4.2. Forced Swim Test

Depressive-like behavior was approached using the FST [44] as previously described [45–47]. Briefly, mice (n=5 of each experimental condition) were individually placed for 6 min in a glass cylinder filled with 17cm of 32±1 °C water. The latency to the first immobilization as well as the total immobility duration were scored.

#### 2.4.3. Novelty-Suppressed Feeding Task

This behavioral task has been proven useful to evaluate hyponeophagia and the emergence of negative valence behaviors in rodents [48]. Literature considers the inhibition of feeding produced by the novel environment an anxious-like behavior, as it reflects the appetitive component of incentive motivation [49]. Besides being sensitive to anxiolytics, the NSF task is also known to respond to chronic, but not acute, antidepressant treatment, supporting the predictive validity of this paradigm to evaluate comorbid anxio-depressive-like phenotypes in rodents [48,50,51]. As previously described by our team [21], the NSF task took place in an open-field apparatus in the center of which two grain-based pellets were located on a piece of filter paper. The behavior of 24 h food-deprived animals (VEH+H_2_O, n=5; VEH+PEA, n=5; CORT+H_2_O, n=10 and CORT+PEA, n=10) was recorded and their latency to start feeding used as an index of anxio-depressive-like behavior. The task lasted a maximum of 15 min. Animals’ feeding drive was controlled after the completion of the task through measuring the food consumed during 5 additional min [52].

### 2.5. Animals sacrifice and tissue sampling

Sixty-to-eighty min after the NSF task (time course corresponding to the peak response of cFos protein [53,54]), animals (n=5 for each experimental condition) were sacrificed for evaluation of cFos expressing cells in order to examine neuronal activation related to the completion of the NSF task.

The sacrifice and brain sampling procedures were equivalent to those previously described by our team [21]. Following post-fixation and cryoprotection, the olfactory bulbs (OB) were resected and embedded in Tissue-Tek® O.C.T (*Sakura Finetek*, USA, Inc.). Tissue samples were then frozen by immersion in isopentane (2-méthylbutane, *Roth®*, Karlsruhe, Germany) and cut in coronal serial sections (30 μm-thick for the brain using a cryotome, and 10 μm-thick for the OB using a cryostat).

### 2.6. Immunohistochemistry

Several brain sections were selected to study the following regions of interest (ROI) relevant for olfactory and emotional processing: OB at (5.345-3.745 mm) anterior to Bregma (aB), according to the Allen Mouse Brain Atlas, 2008, available at Allen Brain Atlas website https://mouse.brain-map.org/experiment/thumbnails/100142143?image_type=atlas, 2022, [55]; anterior and posterior piriform cortex (aPirCx and pPirCx) at (1.94-1.34 mm and 1.18-0.26 mm) aB; nucleus accumbens (NAc), including its core (NAcC) and shell (NAcS) subregions at (1.70-0.86 mm) aB; anterior insular cortex (aIC) at (1.54-0.50 mm) aB; amygdala complex, including the basolateral (BLA), central (CeA) amygdala and anterior and posterolateral cortical amygdaloid areas (ACo+PLCo) at (0.94-1.70 mm) posterior to Bregma (pB); hippocampus (Hip) at (0.94-1.94 mm) pB; parasubthalamic nucleus (PSTN) at (2.06–2.54 mm) pB; and lateral entorhinal cortex (LEC) at (2.18-3.08 mm) pB (according to Franklin and Paxinos, 2001, [56]). The immunostaining protocol was performed as previously described [21]. Briefly, after being exposed to an antigen retrieval method, slide mounted sections were incubated overnight at room temperature with a rabbit anti-cFos monoclonal primary antibody (9F6, *Cell Signaling*, 1:8000), diluted in 0.1M phosphate buffer at pH 7.4, containing 0.3% Triton X-100, 1% bovine serum albumin and 0.01% sodium azide (blocking buffer used as diluent for primary and secondary antibodies). Sections were then washed and incubated overnight with the secondary biotinylated antibody (BA-1000, goat anti-rabbit IgG, *Vector Laboratories*, 1:1000). Subsequently, classical avidin-biotin-horseradish peroxidase procedure (Vectastain Elite ABC Kit, *Vector Laboratories*) was used. The peroxidase complex was visualized by exposing all tissue sections to a chromogen solution containing 0.04% 3,3’-Diaminobenzidine tetrahydrochloride chromogen (DAB, *Sigma Aldrich*®, France), 0.006% hydrogen peroxide (*Sigma Aldrich*®, France), and 0.04 % Nickel (II) chloride (*Roth*®, Germany). Finally, sections were dehydrated through successive baths of ascending ethanol concentrations, cleared with xylene, and then coverslipped.

Stained tissue sections were visualized using 4x or 10x objectives (for brain and OB sections, respectively) of an Olympus microscope Bx51 equipped with a digital camera Olympus DP50. Images analysis and cFos-labelled cell quantification within each ROI were performed using *ImageJ* software (*National Institutes of Health*, Bethesda, MD, USA [57]) according to the protocol previously described [58]. While a frame delimiting the entire area of each brain ROI was manually traced, a sampling window (200 μm × 200 μm) placed always in the same position (to include the mitral and granule cell layers) within the selected area was used for OB sections. Coronal OB sections were subdivided into four quarters corresponding to the dorsolateral (DL), dorsomedial (DM), ventrolateral (VL), and ventromedial (VM) areas [59]. The number of cFos-immunoreactive nuclei in each quadrant was counted according to the selected area (cFos+/mm^2^) and the data obtained were then cumulated to reflect a single value of cell density for each OB section.

Due to technical issues, some regions could not be included in the final analysis, so that final samples sizes for cFos quantification were: BO n=20, aPirCx n=14, pPirCx n=18, aIC n=19, aIC n=17, NAcC n=16, NAcS n=16, BLA n=18, CeA n=18, ACo+PLCo n=18, Hip n=19, PSTN n=19, LEC n=17.

### 2.7. Data and statistical analyses

Data are presented as means ± SEM and statistical analyses were conducted using STATISTICA 10 (*Statsoft*, Palo Alto, United States). GraphPad Prism 9 software (*GraphPad Inc*., San Diego, United States) was used for designing the figures. Assumptions for parametric analysis were verified prior to the analyses: normality of distribution with Shapiro-Wilk and homogeneity of variance with Levene’s tests. Since datasets did not meet assumptions for parametric analyses, Mann-Whitney U (MWU) tests were used for comparing behavioral and immunohistological scores between experimental conditions. Survival curves for NSF scores were compared using the Long-rank Mantel-Cox (MCox) test. The relationship between NSF behavioral scores and cFos expression in the various brain structures under investigation were analyzed using Pearson correlations. Pearson *r*-values from interregional c-Fos expression data within experimental conditions were correlated using cross-correlation matrices. For all analyses, the significance level was set up at p < 0.05, and analyses with p ≤ 0.1 were described as trends.

## 3. Results

### 3.1. Behavioral evaluation

The therapeutic potential of chronic PEA inhalation to reverse the pathological anxio-depressive-like phenotype induced by chronic GC administration in female mice was addressed using various behavioral paradigms. Respectively, the OF test and the FST allowed measuring anxiety- and depressive-like behaviors, while the NSF task allowed evaluating depression-related anxiety.

Contrary to expected, the emergence of anxious-like behavior resulting from a chronic exposure to CORT was not evidenced in this study through the OF evaluation. Female mice treated with CORT spent a similar relative time in the center of the arena (all groups n=10; CORT-H_2_O: mean duration (s) ± SEM = 15.27±2.90) than VEH-treated mice (VEH-H_2_O: 13.10±2.76) [MWU, Z=-0.42, p=0.68]. However, a significant effect of the treatment on the animals’ locomotor activity was evidenced [Z=3.33, p<0.001], with CORT-treated individuals travelling a shorter distance in the 5 min of testing (mean travelled distance (m) ± SEM = 26.28±2.43) than control individuals (41.27±1.62). Most importantly, when evaluating the therapeutic effect of PEA in CORT-treated mice, the results show that the olfactory inhalation protocol did not significantly impact the time animals spent in the center of the arena (CORT-PEA: 21.03±5.04) [Z=-0.64, p=0.52], but tended to improve their locomotor activity (CORT-PEA: 34.72±3.05) [Z=-1.889, p=0.06]. The inhalation of PEA *per se* had no significant behavioral effect on VEH-treated animals, neither on the time in central arena (VEH-PEA: 15.39±2.34) [Z=-0.53, p=0.60], nor on the locomotor activity of mice (VEH-PEA: 48.43±3.52) [Z=-1.51, p=0.13]. Indeed, further comparisons suggest that the potential therapeutic effect of PEA inhalation on locomotor activity is restricted to the existence of pathological traits, as it is the case in CORT-treated animals [VEH-H_2_O vs CORT-PEA: Z=2.04, p<0.05]. This synergetic treatment/odor effect was not evidenced in the case of the relative time animals spent in the center of the arena during the OF evaluation [Z=-1.13, p=0.26]. See **Fig. 2A&B**.

**Figure 2.**
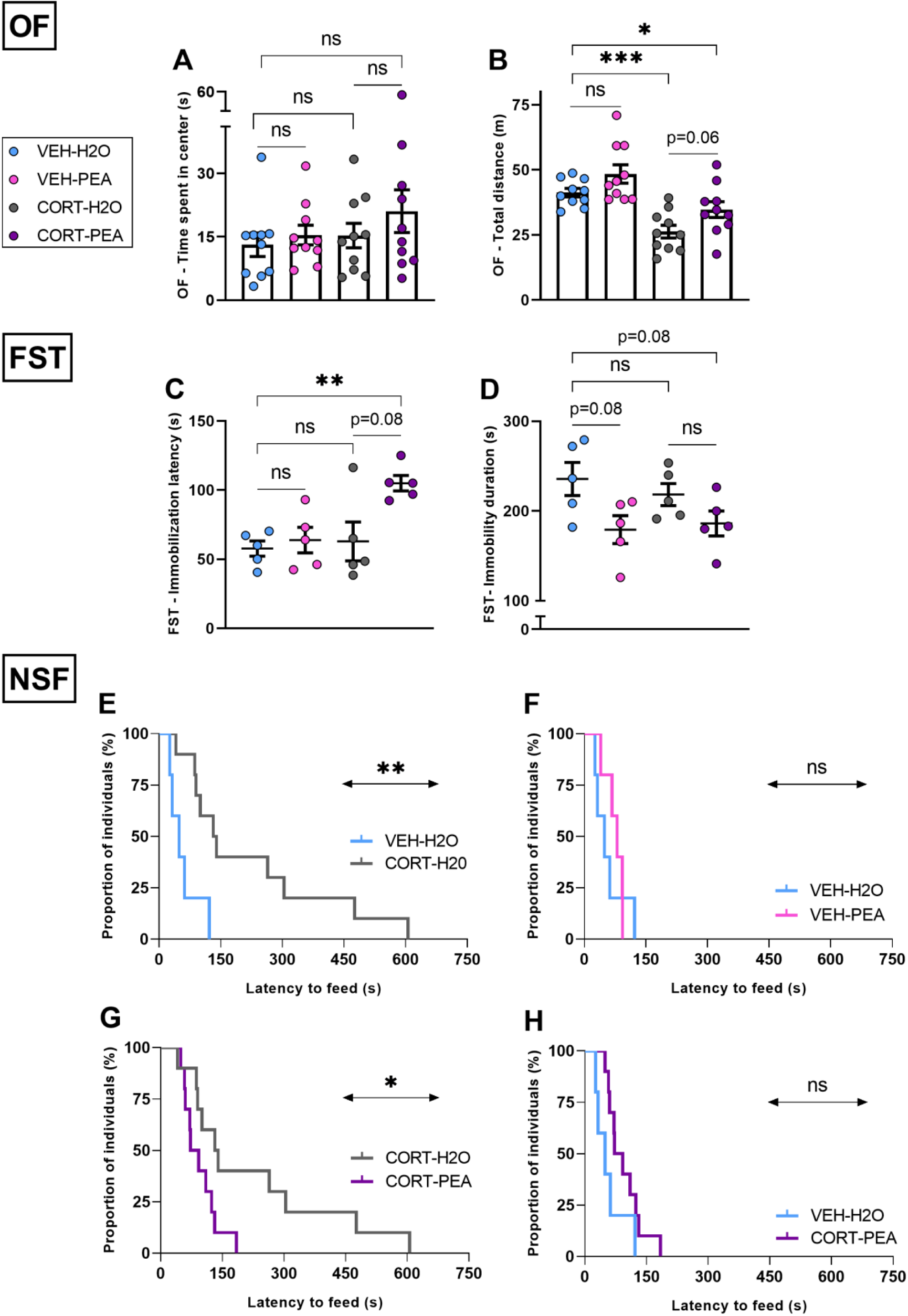
Effect of chronic PEA inhalation on CORT-induced behavioral changes. **(A)** The anxiety-like behavior assessed in the OF was not affected neither by the chronic CORT treatment nor the PEA inhalation. **(B)** The chronic odor inhalation protocol tended to improve (p=0.06) the reduced locomotor activity observed in CORT-treated mice (***, p<0.001). A synergetic treatment/odor effect was revealed in CORT animals when compared to the physiological condition (*, p<0.05). **(C, D)** In the FST, CORT had no effect on the latency to first immobilization or the immobility duration, but chronic PEA inhalation positively influenced the animals’ coping ability by increasing the latency to the first immobility event (**, p<0.01). **(E)** In the NSF task, CORT administration induced an anxio-depressive-like phenotype by increasing the animals’ latency to feed (**, p<0.01). **(F, G, H)** Although no odor-evoked behavioral effect was observed in VEH-treated mice, chronic PEA reversed the pathological phenotype seen in CORT-treated animals (*, p<0.05) to a similar level to that revealed in the VEH-treated group. Values plotted are mean ± SEM (n=5–10 per group). Not significant, ns.

In order to investigate whether chronic PEA inhalation had a positive effect on the animals’ coping ability when facing a stressful situation, mice were tested in the FST (all groups, n=5). Unexpectedly, mice chronically treated with CORT were found to behave similarly to VEH-treated mice in this paradigm: they reached the first immobilization after a similar latency(VEH-H_2_O: mean immobilization latency (s) ± SEM = 57.63±5.50; CORT-H_2_O: 62.91±14.05) [Z=0.52, p=0.60], and adopted an immobile posture for similar durations within the 6 min of testing (VEH-H_2_O: 235.67±18.47; CORT-H_2_O: 218.28±12.29) [Z=0.52, p=0.60]. However, a trend was evidenced for the influence of PEA exposure on animals’ behavior. After chronic CORT administration, PEA inhalation tended to increase the latency of the first immobilization (CORT-PEA: 104.97±5.58) [Z=-1.78, p=0.08], but did not influence the total immobility duration during the FST (CORT-PEA: 186.15±13.92) [Z=1.57, p=0.12]. The PEA inhalation protocol had no significant effect on the coping strategy of animals when they were VEH-treated, but tended to decrease their total immobility duration (VEH-PEA, immobilization latency: 63.76±9.25; immobility duration: 179.19±15.50) [immobilization latency: Z=-0.52, p=0.60; immobility duration: Z=1.78, p=0.08]. Most interestingly, a synergetic CORT/PEA effect was revealed concerning the latency of the first immobilization [VEH-H_2_O vs CORT-PEA: Z=-2.61, p<0.01], in line with the OF results. Although this effect did not reach significance for the total immobility duration, CORT-treated animals exposed to chronic PEA tended to more actively cope with the stressful situation [Z=1.78, p=0.08]. See **Fig. 2C&D**.

The NSF task allows evaluating co-existing anxio-depressive behaviors, since mice hyponeophagia refers to both the anxiety-related dimension of exposing themselves in a novel environment, and to the motivational dimension of behavioral initiation frequently curtailed in depressive-like phenotypes. As expected, 8 weeks of CORT administration significantly prolonged the time mice required to start eating from the available food (CORT-H_2_O, n=10: mean latency to eat (s) ± SEM = 224.00±59.49) compared with animals from the control group (VEH-H_2_O, n=5: 58.20±17.17) [MCox, X^2^=7.71, p<0.01] (**Fig. 2E**). Chronic exposure to CORT, therefore induced a typical negative valence behavior in the NSF task, in line with previous studies [13,18,21,41]. Concerning the PEA effects on animals’ behavior, the odor inhalation had not significant effect *per se* when administered to control animals (VEH-PEA, n=5: 74.40±9.81) [X^2^=0.20, p=0.65] (**Fig. 2F**), but significantly improved the anxio-depressive-like phenotype in CORT-treated animals (CORT-PEA, n=10: 95.10±13.38) [X^2^=4.93, p<0.05] (**Fig. 2G**). Indeed, after chronic PEA inhalation CORT-treated mice tended to behave as control animals did in the NSF task [X^2^=3.03, p=0.08] (**Fig. 2H**), supporting the therapeutic potential of the odor inhalation protocol to reverse the CORT-induced anxio-depressive-like phenotype.

### 3.2. Neurobiological approach

In this work, the potential therapeutic effect of chronic PEA inhalation on CORT-induced anxio-depressive states was for the first time approached by studying its action on neural activation. Thus, in order to investigate whether the behavioral improvement observed in CORT-PEA individuals during the NSF task could result from different patterns of activation, we selected a set of brain regions based on their role in neuronal circuitry involved in olfactory processing (OB, aPirCx, pPirCx, ACo and PLCo, LEC, Hip) and feeding behavior control (aIC, BLA, CeA, PSTN). The NAc was also included in the network analysis to approximate the possible implication of the reward system on the beneficial effect of chronic PEA.

The density of cFos expressing cells evaluated after the NSF task, and the comparisons between experimental conditions for each brain region are presented in **Table 1**. Of most importance for the present study, the density of cFos-labelled cells was significantly decreased in the OB of mice chronically treated with CORT compared to control animals [MWU, Z=2.6, p<0.01]. The enhancing effect of chronic PEA inhalation on OB neural activation was also revealed when comparing CORT-H_2_O and CORT-PEA experimental conditions [Z=-2.6, p<0.01], and confirmed in the comparison between CORT-PEA and the control condition, where no significant difference was evidenced [Z=-1.1, p=0.25] (see **Fig. 3A-F**). Therefore, these results suggest that sustained CORT administration impacts the first stage of olfactory information processing, and that chronic PEA inhalation is effective in reversing this effect. In addition, the density of cFos-labelled cells in the OB was found to negatively correlate with the latency to start feeding in the NSF task [OB: r=-0.5073, p<0.05; other regions: p>0.05], supporting our findings and revealing the key role of olfactory processing in this behavioral test (see **Fig. 3G**).

**Table 1.** Data of cFos expression after Novelty-Suppressed Feeding task evaluation in the selected brain regions of interest, and statistical comparison between experimental conditions. *, p<0.05; **, p<0.01.

| ROI | cFos expression (mean cFos+ cells/mm<sup>2</sup> ± SEM) |  |  |  |
| --- | --- | --- | --- | --- |
|  | VEH-H<sub>2</sub>O | VEH-PEA | CORT-H<sub>2</sub>O | CORT-PEA |
| OB | 62.72±8.11 | 78.62±9.09 | 25.56±2.00 | 74.70±9.57 |
| aPirCx | 856.98±123.77 | 992.49±44.13 | 1046.31±158.65 | 1226.75±93.95 |
| pPirCx | 847.13±83.13 | 1067.05±81.12 | 998.14±117.66 | 975.05±65.27 |
| NAcC | 370.11±48.57 | 415.68±40.72 | 280.43±53.35 | 432.00±34.69 |
| NAcS | 453.53±57.01 | 500.58±28.07 | 512.24±94.38 | 460.47±9.12 |
| aIC | 263.53±27.69 | 374.20±56.67 | 385.82±56.93 | 453.25±59.32 |
| BLA | 259.23±17.76 | 358.61±21.52 | 294.52±59.02 | 358.79±44.62 |
| CeA | 260.81±20.01 | 360.24±39.51 | 347.13±62.77 | 340.40±17.58 |
| ACo+PLCo | 371.21±63.82 | 422.31±46.38 | 591.38±39.76 | 733.83±94.17 |
| Hip | 184.08±18.61 | 202.39±9.33 | 252.56±30.50 | 264.09±6.74 |
| PSTN | 221.62±21.20 | 254.21±34.78 | 226.80±74.40 | 383.24±28.91 |
| LEC | 264.67±27.56 | 470.99±24.24 | 324.82±30.22 | 346.67±25.75 |
| Statistical difference (MWU), *p* value |  |  |  |  |
|  | VEH-H<sub>2</sub>O vs VEH-PEA | VEH-H<sub>2</sub>O vs CORT-H<sub>2</sub>O | CORT-H<sub>2</sub>O vs CORT-PEA | VEH-H<sub>2</sub>O vs CORT-PEA |
| OB | 0.17 | ** | ** | 0.25 |
| aPirCx | 0.25 | 0.33 | 0.25 | 0.050 |
| pPirCx | 0.17 | 0.46 | 0.77 | 0.33 |
| NAcC | 0.46 | 0.33 | 0.050 | 0.56 |
| NAcS | 0.46 | 0.46 | 0.33 | 0.25 |
| aIC | 0.12 | 0.12 | 0.62 | * |
| BLA | * | 1 | 0.39 | 0.050 |
| CeA | * | 0.33 | 0.77 | * |
| ACo+PLCo | 0.46 | 0.050 | 0.25 | 0.050 |
| Hip | 0.35 | 0.086 | 0.81 | ** |
| PSTN | 0.25 | 0.62 | 0.086 | ** |
| LEC | ** | 0.12 | 0.81 | 0.050 |

**Figure 3.**
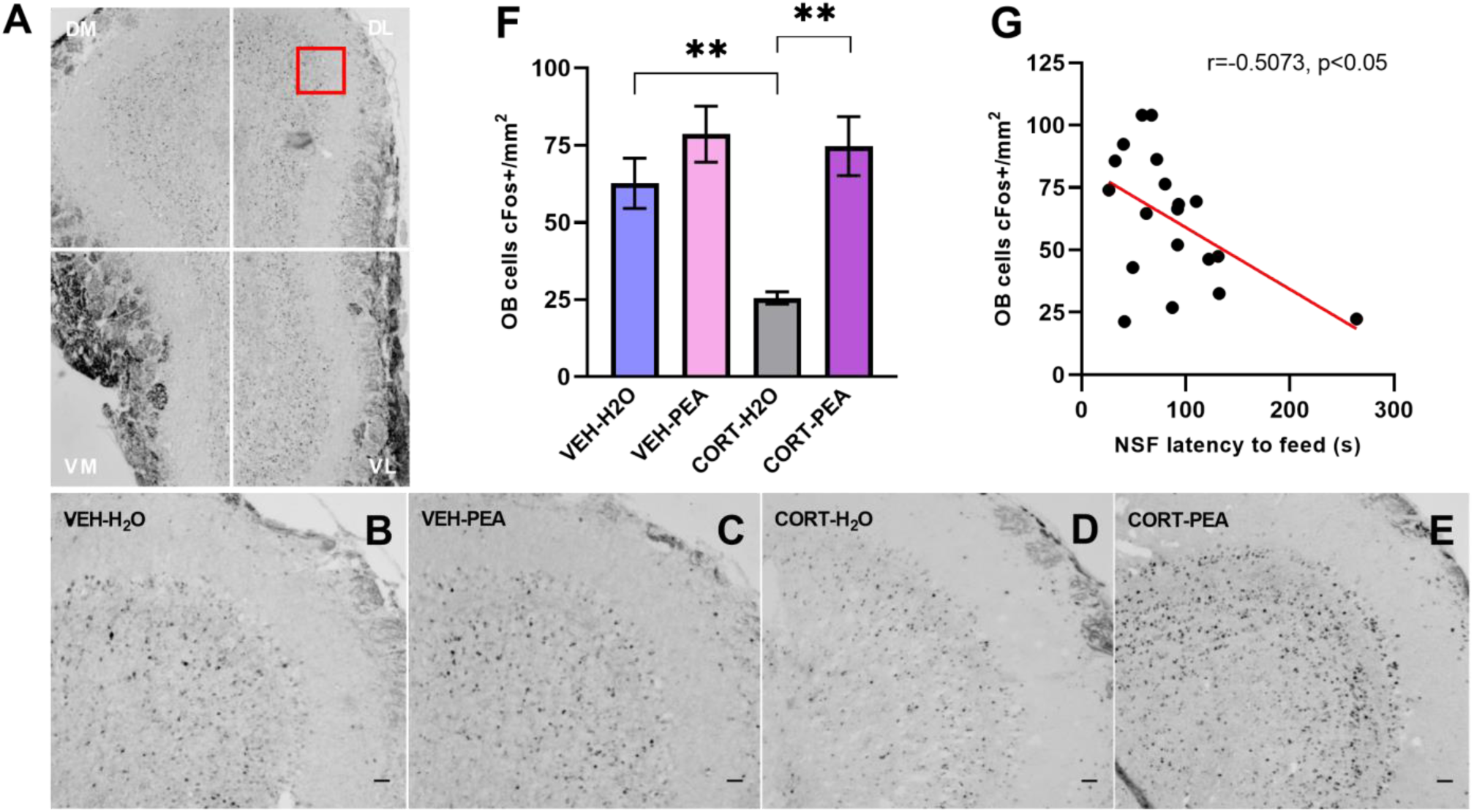
cFos expression in the olfactory bulb (OB) following the NSF task. **(A)** Photomicrograph of a representative coronal OB section, indicating the four quadrants in which cFos-immunoreactive cells in the mitral and granule cell layers were quantified (DM, dorsomedial; DL, dorsolateral; VM, ventromedial; and VL, ventrolateral areas of the OB). Sampling window in red: 200μm x 200μm. **(B-E)** Representative images of cFos-labelled nuclei in the DM quadrant of the OB (scale bar: 100 μm). **(F)** Histogram representing the density of cFos-immunopositive cells (number of c-Fos labelled cells/mm^2^) in the OB for each experimental condition (n=5 per group). Values plotted are mean ± SEM. CORT administration significantly decreased the density of cFos-labelled cells in the OB (**, p<0.01). This deleterious effect was reversed by chronic PEA inhalation (**, p<0.01). **(G)** A negative correlation was found between behavioral performances in the NSF task and OB cFos-labelled cell densities (r=-0.5073, p<0.05), indicating the importance of the main olfactory system in feeding behavioral control.

Finally, in order to infer interactions and to approximate the strength of coordinated activity changes among brain regions, correlation coefficients across experimental conditions were computed using the cFos expression data set. The generated cross-correlation matrices (**Fig. 4A-D**) show changes in network density upon CORT and PEA exposures. More specifically, chronic CORT administration induced a reorganization of the coordinated activity, reducing the synchronization of the main actors of the olfactory processing and feeding behavior control circuitries (**Fig. 4A&B**). The action of chronic PEA inhalation on neural activation was evidenced in VEH-treated animals, even if its effect did not translate in a NSF behavioral output. Indeed, the cross-correlation matrix of non-distressed animals exposed to the odor displays a significant desynchronization, or even an opposite pattern of coordination, between the activity of both circuitries (**Fig. 4C**). Finally, chronic PEA inhalation modified the pattern of coordinated neural activity of CORT-treated animals, partially recovering the synchronization of the involved circuitries observed in control individuals (**Fig. 4D**). Therefore, this odor-induced resynchronization of the neural network might mediate the NSF behavioral output, i.e. VEH-H_2_O and CORT-PEA individuals behaving similarly in this paradigm. Interestingly, an inversion of the PSTN coordinated activity pattern as compared to the physiological condition is observed in CORT-PEA individuals. This hypothalamic structure, which acts in a coordinated manner with the other brain regions studied in VEH-H_2_O animals, appears strongly anticorrelated with the majority of neural actors in the CORT-PEA condition. Of particular interest, while displaying a co-depression pattern in VEH-H_2_O animals, the PSTN reveals a co-activation pattern with the CeA in CORT-treated animals. Most importantly, the PEA inhalation seems to normalize the initial functional connectivity between PSTN and CeA in CORT-PEA mice. These findings therefore suggest a particular involvement of the PSTN in the reversion of the anxio-depressive-like phenotype induced by chronic CORT by regulating the consummatory behavior controlling the food intake. In addition to computing interregional correlations, general changes in density network were studied from the mean *r* values obtained for each experimental condition. As shown in **Fig. 4E**, a tendency to a higher functional connectivity between the selected brain regions was observed upon chronic CORT treatment (mean r ± SEM, VEH-H_2_O: 0.32±0.04; CORT-H_2_O: 0.42±0.04) [*t*_286_=-1.8, p=0.07], which was reversed after inhalation of PEA (CORT-PEA: 0.32±0.04) [*t*_286_=0, p=1]. Interestingly, PEA chronic exposure in VEH-treated animals significantly impacted functional network connectivity in relation to the physiological condition (VEH-PEA: 0.03±0.04) [*t*_286_=4.7, p<0.001].

**Figure 4.**
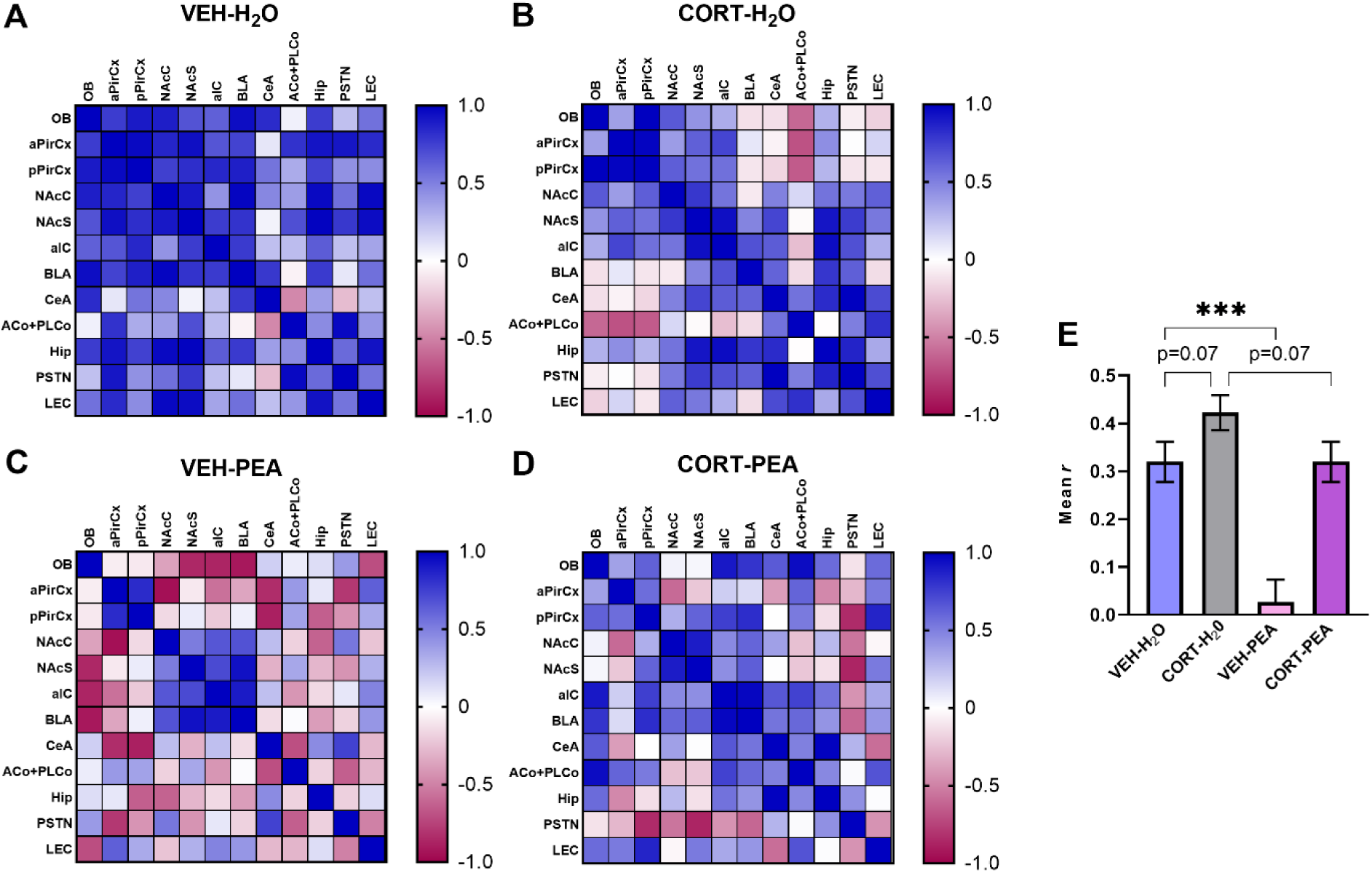
Cross-correlation matrices of functional connectivity of the brain regions of interest. **(A, B)** CORT administration altered the coordinated activity among the selected brain regions involved in olfactory processing and feeding behavior control circuitries. **(C)** Odor-evoked changes in overall network activity were revealed in VEH-treated mice, confirming the effect of chronic PEA inhalation on neural functionality. **(D)** Chronic PEA inhalation led to a partial recovery of the functional connectivity, with the exception of the opposite activity pattern of the PSTN, suggesting its implication in the effect of PEA on feeding behavior. Values close to 0 (white) indicate absence of linear correlation between regions’ activity; values of 1 (blue) indicate a positive linear correlation; and values of -1 (red) indicate a negative linear correlation. **(E) Mean *r* calculated from the correlation matrices of each experimental condition**. Changes in network density were observed after both CORT and PEA exposure (***, p<0.001).

These findings further support the occurring functional connectivity changes upon sustained GC treatment and PEA odor exposure.

## 4. Discussion

In this investigation we evaluated the potential therapeutic action of prolonged PEA inhalation on mood-related symptomatology in chronically CORT-treated female mice. Overall, our behavioral approach revealed more of an antidepressant-like profile of PEA inhalation rather than an anxiolytic-like effect. Specifically, a positive impact of long-term PEA exposure was shown on locomotor function, coping ability and initiation of feeding behavior, assessed respectively in the OF, FST and NSF paradigms. Moreover, our results suggest that the behavioral effects of chronic PEA inhalation in CORT-treated mice might be mediated via the olfactory system which is reciprocally connected with neural circuits engaged in emotional and feeding behaviors.

An increasing body of evidence confirms the deleterious effect of chronic CORT administration in male mice [13,14,21,45,60–63], but data concerning the behavioral and neural changes occurring in female mice are scarce. Few available studies [41,64–66] suggest that female mice are less sensitive to CORT treatment compared to their male counterparts, but still display a higher emotionality score in contrast to control animals. In agreement with these findings, the present study did not reveal neither a strong anxiogenic-nor a marked depressant-like effect of chronic CORT administration in female C57BL/6JRJ mice, when they were evaluated in the OF and FST, respectively. This may probably reflect the lower sensitivity of female mice to the chronic CORT treatment and the associated pathological phenotype. However, a significant alteration of the overall motor activity was evidenced in the OF paradigm, which could be associated with the emergence of motor retardation leading to deleterious consequences on physical activities. Our behavioral data is in line with other preclinical studies demonstrating decreased locomotor activity in CORT-treated male mice [15,18]. The hampering effect of HPA axis dysfunction on motor performance was also observed in male mice in our previous study [45], thus evoking the clinical phenotype reported among depressed patients [67]. Interestingly, our chronic PEA inhalation protocol tended to restore the CORT-induced motor deficit to a more physiological level to that observed in VEH-treated animals, suggesting a stimulant effect in female mice. Such activating action on locomotion was demonstrated in a previous study from our team using naive female OF1 mice exposed to acute PEA odor during the OF [39]. Although CORT-treated animals did not show a depressive-like phenotype in the FST, the chronic PEA inhalation was revealed to play a role in helping to actively cope with the stressful situation. In fact, the latency to the first immobilization increased to a much higher level than that observed in VEH-treated mice, which further supports the stimulant effect of PEA observed in the OF. Consistent with previous findings in male [13,18,21] and female mice [41], our experimental set-up further showed that CORT-treated animals displayed a significantly longer latency to feed in the NSF task, indicating additional disturbances in motivation-guided feeding behavior. Besides being sensitive to anxiolytic drugs, the NSF task is also known to respond to chronic, but not acute, antidepressant treatments [48]. Thus, it is considered to be a reliable behavioral paradigm with predictive validity for investigating the therapeutic action of antidepressants that requires several weeks to emerge [50]. Importantly, our 15-day odor inhalation protocol was able to reverse the CORT-induced anxio-depressive-like phenotype in this conflict-based paradigm, strengthening the assumption that chronic PEA displays a similar therapeutic profile as monoaminergic antidepressants in terms of pharmacokinetics [50,68]. Overall, our behavioral results suggest that the beneficial effect of PEA is restricted to the existence of pathological traits since it is devoid of such outcome in VEH-treated animals. Thus, the therapeutic effect of chronic PEA inhalation becomes robust after prolonged CORT-exposure in female mice. A similar combined effect was previously reported [69], demonstrating an anti-stress activity of chronic oral administration of the hydroalcoholic extract of *Rosa gallica officinalis* in male mice subjected to repeated restraint stress protocol. Moreover, it has been shown that naive male C57BL/6 mice are unresponsive to chronic fluoxetine, a selective serotonin re-uptake inhibitor used in the treatment of depression, but the CORT-induced activation of stress circuitry is sufficient to reveal its therapeutic action in this typically unresponsive strain [70].

It is noteworthy that in the present study we used female rather than male mice in order to examine the effect of chronic olfactory PEA exposure on behavioral emotionality. This choice was guided by the evidence of sexual dimorphism in the human main olfactory bulb, with females displaying higher number of neuronal and glial cells than males [71], probably leading to a greater sensitivity towards olfactory cues, as observed in women [72]. Such sexual difference was also demonstrated in adult mice, with a more robust primary sensory odor coding occurring in female compared to male mice [73]. To our knowledge, no study to date has investigated the effects of long-term PEA inhalation on emotional states in chronically distressed female mice. Indeed, most animal studies focusing on rose odor essential oil effects, or its active constituents, have been carried out in naive animals during or immediately after acute odor inhalation [35,39,40]. However, when studying the therapeutic action of aromatic compounds on general well-being, it must be taken into account the fact that in clinical trials aromatherapy is rather prescribed over a long period of time, i.e. in a chronically administrated manner. The administration of PEA in the present study was performed by inhalation, which reflects the usual aromatherapeutic use by vaporization of essential oils in humans [74]. Furthermore, inhalation allows avoiding the additional stress associated with intraperitoneal injection, the delivery method used in the majority of preclinical studies [34,38,75,76]. Thus, delivery by inhalation allows the simultaneous absorption of volatile aromatic compounds through the broncho-pulmonary system and the olfactory system. The first, allows the passage into the systemic blood circulation and then to the central nervous system by bypassing the blood-brain barrier. The second, importantly, displays strong anatomical connections with limbic regions known to exert a direct action on emotion [77]. Interestingly, it has been suggested that olfaction and depression display intricate relationships, since sensory deprivation often leads to depressive states, and reciprocally depressed patients frequently show olfactory impairments [78,79].

To further elucidate neural correlates of our behavioral results, cFos immunostaining was carried out to map the neural activity associated with NSF performance. Importantly, a significant correlation was found between the NSF task results and OB cFos-labelled cell densities. Our data revealed a significant deleterious effect of prolonged CORT exposure on OB activity, probably leading to the anxio-depressive-like phenotype observed in the NSF task. This is in accordance with already published data demonstrating that chronic CORT-based behavioral emotionality is accompanied by olfactory deficits in mice [52], as well as olfactory deficits can translate to chronic distress-induced behavioral changes [80]. It has been shown that these behavioral and functional alterations are further associated with a pronounced reduction in hippocampal and OB neurogenesis, but reversed by prolonged antidepressant treatment with fluoxetine [52]. Similarly, in the present study, we revealed that both anxio-depressive-like phenotype and decreased OB activity were reversed by chronic PEA inhalation, suggesting that the therapeutic action of PEA is mainly exerted via the olfactory pathway. Indeed, PEA is known to possess no intranasal trigeminal properties and to activate selectively the olfactory system [81,82]. Again, this beneficial effect on OB neural activity was only behaviorally translated in chronically distressed mice. No such changes were observed in VEH-treated animals, although our network analyses indicate that PEA-exposed control animals respond to the odorant, with significant changes occurring in BLA, CeA and LEC regions. The activation of these brain areas might therefore reflect the process of memory recall of the reinforcing and motivational properties of the chronically presented PEA odor. The amygdala and the entorhinal cortex receive direct bulbar projections, thus participating actively in odor information processing [83]. Moreover, it has been shown that the amygdala plays a key role in encoding the emotional salience of the olfactory stimulus [84]. As part of the limbic corticostriatal circuitry, it is also known to be implicated in motivational-based appetitive behaviors [21,85]. From its side, the entorhinal cortex is reciprocally connected to the amygdala and hippocampus, and several studies have pointed out its involvement in odor memory formation [86,87]. Compared to the physiological condition, CORT-mice subjected to chronic PEA inhalation showed significant modifications of neural activity among a set of brain regions, namely aIC, CeA, PSTN and Hip. These particular structures are part of a complex network that has been shown to be involved in feeding behavior [88–90], with a key role of PSTN acting as a refraining signal through descending GABAergic projections originating from the CeA [90,91]. Furthermore, it has been described that the PSTN also receives olfactory inputs, mainly from the anterior part of both basomedial and cortical amygdalar nuclei which are densely innervated by the piriform cortex [92–94]. It is therefore conceivable that CORT-induced prolonged distress reinforce the anorexic action of this basal ganglia-like network, and that PEA inhalation may act like a silencing agent in order to relieve the inhibition on food intake. It is well documented that olfactory processing involves a large number of strongly interconnected brain regions that incorporate complex overlapping networks [83], thus creating functional interactions between neural circuits controlling emotional and feeding processes [95–98]. Therefore, it cannot be excluded that the OB neural activity changes could originate from a differential action of centrifugal systems [99,100] occurring upon CORT administration and PEA exposure. Nonetheless, further studies using opto- and chemogenetic approaches are needed to clarify the specific functional meanings of the observed changes and unravel the precise mechanisms underlying the beneficial effect of PEA on feeding behavior in chronically distressed animals. In addition, since the positive impact of odors on mood-related behaviors is predominantly determined by their hedonic value, complementary investigations aiming to examine the possible involvement of the reward circuitry in the therapeutic effect of chronic PEA inhalation are ongoing.

## 5. Conclusion

Together, the presented data indicate that the antidepressant-like effect of chronic PEA inhalation in female mice is potentiated under situations of prolonged stress. Although the exact mechanisms of action remain unknown, the present study provides novel insight into the neuronal circuitry that might contribute to the therapeutic effects of long-term PEA treatment on sensory, emotion and motivation-related behaviors.

## Abbreviations

ACo: anterior cortical amygdaloid area
aIC: anterior insular cortex
aPirCx: anterior piriform cortex
BLA: basolateral amygdala
CeA: central amygdala
CORT: corticosterone
EPM: elevated plus-maze test
FST: forced swim test
GC: glucocorticoids
Hip: hippocampus
HPA: hypothalamic-pituitary-adrenal axis
LEC: lateral entorhinal cortex
MWU: Mann- Whitney U test
MCox: log-rank Mantel-Cox test
NAcC: nucleus accumbens core subregion
NAcS: nucleus accumbens shell subregion
NSF: novelty-suppressed feeding task
OB: olfactory bulb
OF: open-field test
PEA: 2-phenylethyl alcohol
PLCo: posterolateral cortical amygdaloid area
pPirCx: posterior piriform cortex
PSTN: parasubthalamic nucleus
ROI: region of interest
SEM: standard error of the mean
VEH: vehicle

## Author Contributions

Conceptualization of the experimental design: B.R., Y.P. and J-L.M. Methodology, investigation and data acquisition: B.R., L.C., S.C., C.H. Software development: P.A. Data analysis and interpretation: B.R. and L.C. Drafting of the original manuscript: B.R., L.C., P-Y.R. and Y.P. Supervision: Y.P., P-Y.R., E.H., J-L.M. All authors critically revised the work and approved the version to be published.

## Acknowledgements

The authors thank the Animal Facilities of Besançon for technical support.

